# A clinical stage LMW-DS drug inhibits cell infection by coronaviruses and modulates reactive cytokine release from microglia

**DOI:** 10.1101/2022.09.07.506919

**Authors:** Ann Logan, Michela Mazzon, Lars Bruce, Nicholas M. Barnes

## Abstract

Most coronaviruses infect animals including bats, birds and mammals, which act as hosts and reservoirs for the viruses, but the viruses can sometimes move host species and infect humans. Coronoviruses were first identified as human pathogens in the 1960s and now there are seven types known to infect humans. Whilst four of these types cause mild-to-moderate respiratory disease, the other three may cause more severe and possibly even fatal disease in vulnerable individuals particularly, with the most recent SARS-CoV-2 pandemic being associated with severe acute respiratory syndrome (SARS) in many infected people. The aim of the present study was to evaluate the potential of a unique low molecular weight dextran sulphate (LMW-DS) clinical stage drug, ILB^®^, to inhibit infection of human cells by the NL63 coronavirus assessed by immunofluorescence of viral particles, and also to see if the drug directly blocked the interaction of the SARS-CoV-2 viral spike protein with the ACE2 receptor. Furthermore, we evaluated if ILB^®^ could modulate the downstream consequences of viral infection including the reactive cytokine release from human microglia induced by various SARS-CoV-2 variant spike proteins. We demonstrated that ILB^®^ blocked ACE2:spike protein interaction and inhibited coronaviral infection. ILB^®^ also attenuated the omicron-induced release of pro-inflammatory cytokines, including TNFα, from human microglia, indicating control of post-viral neuroinflammation. In conclusion, given the safety profile of ILB^®^ established in a number of Phase I and Phase II clinical trials, these results highlight the potential of ILB^®^ to treat patients infected with coronaviruses to both limit infectivity and attenuate the progression to severe disease. There is now an opportunity to translate these findings quickly by the clinical investigation of drug efficacy.

## Introduction

Coronaviruses are a large family of enveloped, positive-stranded RNA pathogenic viruses that cause infectious disease in animals and humans. For example, the highly transmissible SARS-CoV-2 virus causes coronavirus disease (COVID-19) which causes a mild to moderate respiratory disease in most humans, but susceptible individuals can become dangerously ill, developing severe acute respiratory syndrome (SARS) and acute respiratory distress syndrome (ARDS) that can result in organ damage^1^. The global pandemic of COVID-19 disease that has spread since 2019 has had devastating health, social and economic consequences that remain burdensome^2^. By August 2022 that World Health Organisation report 599,825,400 confirmed cases of COVID-19 and 6,469,458 confirmed deaths (https://www.who.int/emergencies/diseases/novel-coronavirus-2019), representing just over 1% of those infected.

There is now a wealth of information published on the SARS-CoV-2 virus that describe the host receptors, virus transmission, virus structure-function relationships, pathophysiology, co-morbidities, immune response, treatment and the most promising vaccines and anti-viral drugs^3,4^. This explosion of knowledge, and in particular understanding of the mechanisms of SARS-CoV-2 entry into host cells through binding of the viral spike (S) protein to a key receptor, angiotensin-converting enzyme 2 (ACE2), has led to the development of a wide range of vaccines and other anti-viral drugs that are now controlling disease spread^5^ to some extent. However, successive waves of disease derived from emerging and re-emerging viral variants will ensure the persistence of this disease in the global population for the foreseeable future. Simultaneously, persistent and often debilitating, sequelae are being increasingly recognized in convalescent individuals, with fatigue, malaise, dyspnea, defects in memory and concentration and a variety of neuropsychiatric syndromes as the major manifestations, and several organ systems can be involved. Many of these post-viral infection adverse consequences can be linked to neuroinflammation. The socio-economic legacy of this so-called ‘long COVID’ or post-viral syndrome is only now beginning to be realised^6,7^ and it is evident it will also present major health and economic challenges.

There is global need for new and improved adjunct anti-viral agents and, in particular, those that can also reduce the symptoms of post-viral syndrome. Here we report the potential of a new class of regenerative medicine, a unique low molecular weight dextran sulphate (LMW-DS) called ILB^®^, to have broad spectrum anti-viral effects on the infection capability and the downstream pathological processes initiated by coronaviruses and, in particular, variants of the SARS-CoV-2 virus.

## Materials and methods

### ILB^®^

ILB^®^ is the sodium salt of LMW-DS containing 16-19% sulphur with an average MW of 5 kDa (International Publication No. WO 2016/076780, ILB^®^ is in Phase II development to treat the neurodegenerative condition ALS^8^). ILB^®^ was supplied as the sodium salt dissolved in 0.9% NaCl at 100 mg/ml concentration.

### Experimental Procedures

#### 1. Antiviral activity and cytotoxicity of ILB^®^ against human coronavirus NL63

The antiviral activity of eight dilutions of ILB^®^ was explored by pre-incubation with LLC-MK2 cells (LLC-MK2 (rhesus monkey kidney-derived cells) (ATCC CCL-7)) for 1h before virus addition. Experimental controls were (a) Uninfected untreated cells, (b) Infected untreated cells or (c) Positive control drug (Remdesivir (Selleckchem S8932 PHR1258)). Human coronavirus NL63 (BEI Resources NR-470) plus ILB^®^ were left on the cells for the entire duration of the experiment (48 h). The cytotoxicity of the same range of concentrations of ILB^®^ was determined by MTT assay.

LLC-MK2 cells were seeded in 2 × 96-well plates (one for inhibition and one for cytotoxicity) in complete media (DMEM from Gibco, Cat. No. 61965026, supplemented with 10% FBS (Gibco, Cat. no. 10500064) and 1X penicillin/streptomycin (Gibco, Cat. no. 15070063) at 4,000 cells/100μl/well. After seeding, the plates were incubated at RT for 5 minutes for even distribution, and then at 37°C, 5% CO_2_ until the following day. Virus stocks were diluted into supplemented media (DMEM from Gibco, Cat. no. 61965026 supplemented with 2% FBS (Gibco, Cat. No. 10500064 and 1X penicillin/streptomycin (Gibco, Cat. No. 15070063), to reach an MOI of 2.0. To calculate MOI, it was estimated that the cell number had doubled to 8,000 cells/well since plating the day before.

Following ILB^®^ dilution and mixing, the cells were preincubated for 1 h at 37°C in air plus 5% CO_2_ with ILB^®^ concentrations at 0.005, 0.014, 0.041, 0.123, 0.370, 1.111 and 10.000 mg/ml or Remdesivir at 0.009, 0.027, 0.082, 0.247, 0.741, 2.222, 6.667 and 20.000 µM. After 1h, media was removed from the cells and replaced with 50 µl of virus or media (uninfected untreated control) immediately followed by 50 µl of the ILB^®^ dilutions at twice the final concentrations, as they become diluted to the final concentrations by an equal volume of virus or media. Plates were then incubated for 48 h at 37°C in air plus 5% CO_2_, and the ILB^®^ and/or the virus remained with the cultured cells for the entire duration of the experiment. After 48 h, the infection plates were washed with PBS, fixed for 30 mins with 4% formaldehyde, washed again with PBS, and stored in PBS at 4**°**C until staining.

For the infectivity readout, cells were immunostained for relevant viral particles. Briefly, any residual formaldehyde was quenched with 50 mM ammonium chloride, after which cells were permeabilised (0.1% Triton X100) and stained with an antibody recognising double-stranded RNA (Caltag Medsystems Ltd, SCI-10010200). The primary antibody was detected with an Alexa-488 conjugate secondary antibody (Life Technologies, A11001), and nuclei were stained with Hoechst. Images were acquired on an CellInsight CX5 high content platform (Thermo Scientific) using a 4X objective, and percentage infection calculated using CellInsight CX5 software (infected cells/total cells x 100).

For a cytotoxicity test, media was removed from the cells and replaced with 50 µl of supplemented media, followed by 50 µl of the diluted formulations or media. After mixing, the plates were incubated for 48 h at 37°C in air plus 5% CO_2_. Cytotoxicity was detected by MTT assay. Briefly, the MTT reagent (Sigma, M5655) was added to the cells for 2 h at 37°C, 5% CO_2_, after which the media was removed and the precipitate solubilised with a mixture of 1:1 Isopropanol:DMSO for 20 minutes. The supernatant was transferred to a clean plate and signal read at 570nm.

Normalised percentages of inhibition were calculated using the following formula:

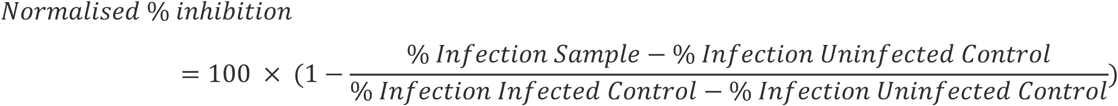

Where appropriate, IC_50_ values were generated by iterative curve fitting according to a Logistic equation using KaleidaGragh software (v5.0; Synergy Software).

Percentages of cytotoxicity were calculated using the following formula:

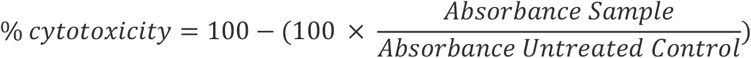

Where appropriate, TC_50_ values were generated by iterative curve fitting according to a Logistic equation using KaleidaGragh software (v5.0; Synergy Software).

#### 2. ILB^®^ inhibition of the interaction between SARS-CoV-2 Spike protein and the ACE2 receptor

The RayBiotech^®^ Life COVID-19 Spike-ACE2 binding assay kit (Catalogue Number: CoV-SACE2) was used exactly as described in the manufacturer’s instructions (Ver 1.6) to assess the ability of a range of concentrations of ILB^®^, between 0.3 and 600 µg/ml, to disrupt the interaction between SARS-CoV-2 Spike protein and ACE2.

#### 3. ILB^®^ attenuation of Coronavirus-stimulated cytokine release from human microglia

Peripheral blood mononuclear cells (PBMCs) were isolated from three healthy donors through SepMate density centrifugation (StemCell Technologies, 85450) with Ficoll-Paque PLUS (Cytiva 17-1440-03). Monocytes were purified from the PBMC population using the EasySep Human monocyte enrichment kit (STEMCELL Technologies; 19059), plated in 96-well plates and differentiated into microglia (iMDM) with the addition of cytokines; M-CSF, GM-CSF, NGF-β, CCL2 and IL-34 for 5 days at 37°C.

iMDM were pre-incubated in the absence or presence of ILB^®^ (600 mg/mL) for 30 minutes prior to the addition of recombinant SARS-CoV-2 spike protein (‘original [Wuhan]’ (Acro Biosystems SPN C52H9), ‘Delta’ [B.1.617.2] (Acro Biosystems SPN-C52He), ‘Omicron’ [B.1.1.529] (Acro Biosystems SPD-C522e); (1.0 or 5.0 mg/mL) in the absence or presence of cross-linker (Abcam; ab18184), and culture for a further 6 or 20 h. Positive control stimulation of microglia was also performed by the addition of LPS (100 ng/mL; Invivogen tlrl-b5lps) and, where indicated, BzATP (100 µM) stimulation for the final two hours of the culture period for 6 or 20 h. After 6 and 20 h, the cell culture supernatants were collected and stored at -20°C for subsequent quantification of the levels of IL-1β, IL-6, MCP-1 and TNFα by Luminex (R&D Systems; LXSAHM-04).

## Results

### 1. Antiviral activity and cytotoxicity of ILB^®^ against human coronavirus NL63

Table 1 displays the EC_50_, EC_90_, TC_50_, TC_90_ and Selectivity Index (SI_50_) and SI_90_ for ILB^®^ and the Remdesivir control. Inhibition of human coronavirus NL63 infection was observed for both the test and control formulations, with an EC_50_ of 5.90 mg/ml for ILB^®^ and 1.32 μM for Remdesivir. Cytotoxicity was observed only at the highest concentrations of ILB^®^, with TC_50_ of 56.5 mg/ml. No significant cytotoxicity was observed at any of the concentrations of Remdesivir tested.

**Table 1.**
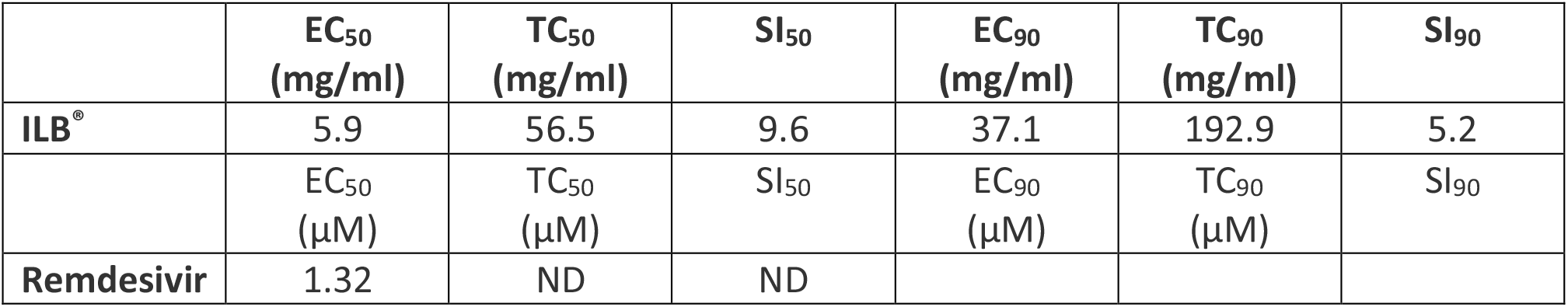
EC_50_, EC_90_, TC_50_, TC_90_, SI_50_ (=TC_50_/EC_50_) and SI_90_ (=TC_90_/EC_90_) values for NL63 against ILB^®^ and Remdesivir. ND: Not determined, due to the inability to extrapolate a curve or the Hill Slope value from the input values.

|  | EC<sub>50</sub><br>(mg/ml) | TC<sub>50</sub><br>(mg/ml) | SI<sub>50</sub> | EC<sub>90</sub><br>(mg/ml) | TC<sub>90</sub><br>(mg/ml) | SI<sub>90</sub> |
| --- | --- | --- | --- | --- | --- | --- |
| ILB<sup>®</sup> | 5.9 | 56.5 | 9.6 | 37.1 | 192.9 | 5.2 |
| | EC<sub>50</sub><br>(μM) | TC<sub>50</sub><br>(μM) | SI<sub>50</sub> | EC<sub>90</sub><br>(μM) | TC<sub>90</sub><br>(μM) | SI<sub>90</sub> |
| Remdesivir | 1.32 | ND | ND |  |  |  |

The graphs in Figure 1 below display the percentage of inhibition of human coronavirus NL63 infection at different ILB^®^ concentrations compared to Remdesivir. Sample values in each plate were normalised to the plate internal controls, where 100% inhibition was derived from the average of the negative control (untreated uninfected) and 0% inhibition was derived from the average of the positive control (untreated infected). The x axes show compound dilutions (mg/ml [ILB^®^] or μM [Remdesivir]). The curves represent the best fit of the logarithm of compound dilution vsersus the normalised percentage of inhibition (variable slope). Cytotoxicity is displayed in grey, with values normalized to the plate internal control (untreated cells, 100% viability). Percentages of infection and cytotoxicity relative to each ILB^®^ concentration are shown in the Appendix as Supplementary Tables S1 and S2 respectively.

**Figure 1.**
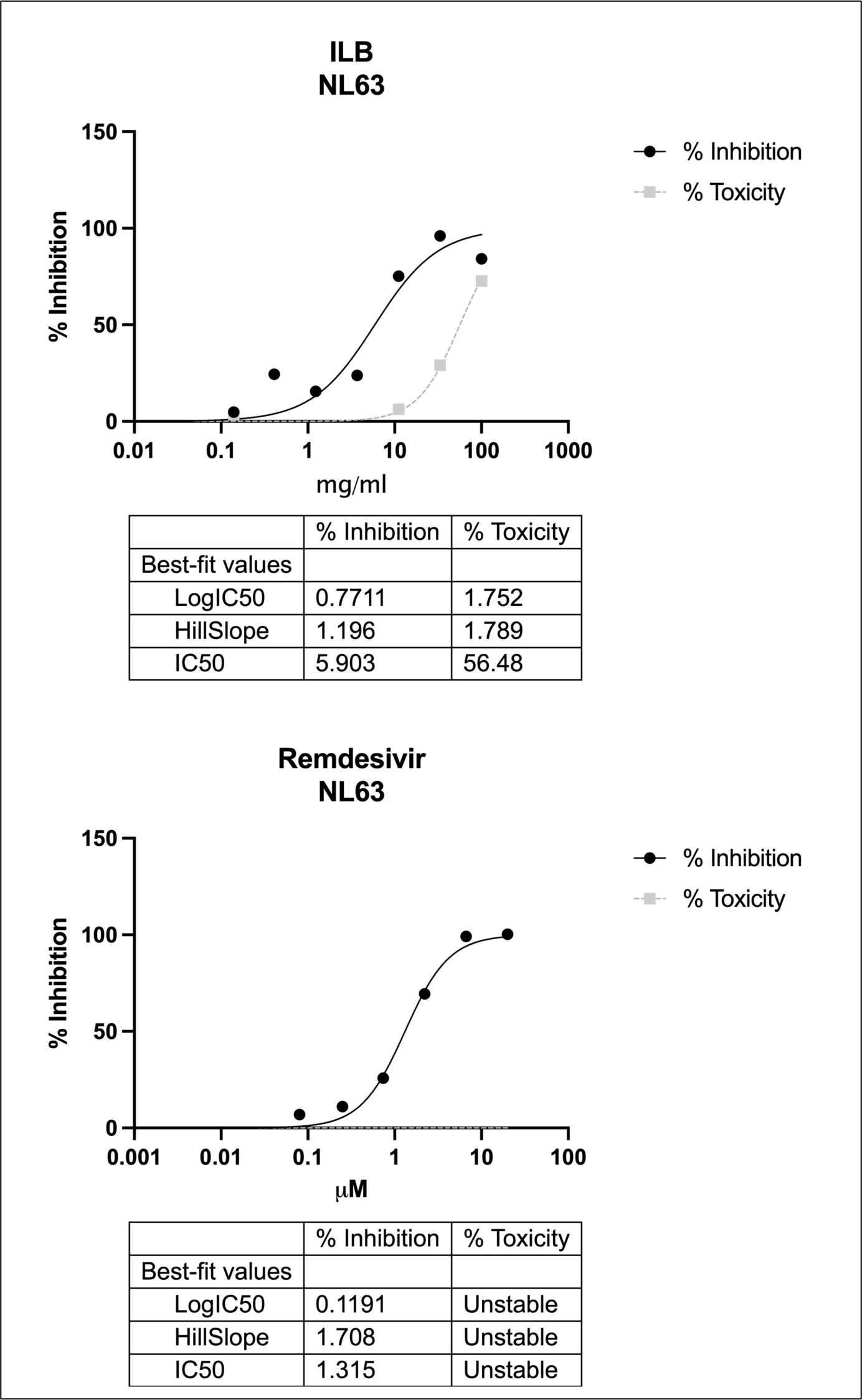
The percentage of inhibition of human coronavirus NL63 infection by ILB^®^ and Remdesivir (black lines) at different compound concentrations. Cytotoxicity is displayed in grey.

### 2. ILB^®^ inhibition of the interaction between SARS-CoV-2 Spike protein and the ACE2 receptor

In each of three independent experiments ILB^®^ reduced the interaction between SARS-CoV-2 spike protein and ACE2 (see representative data in Figure 2).

**Figure 2.**
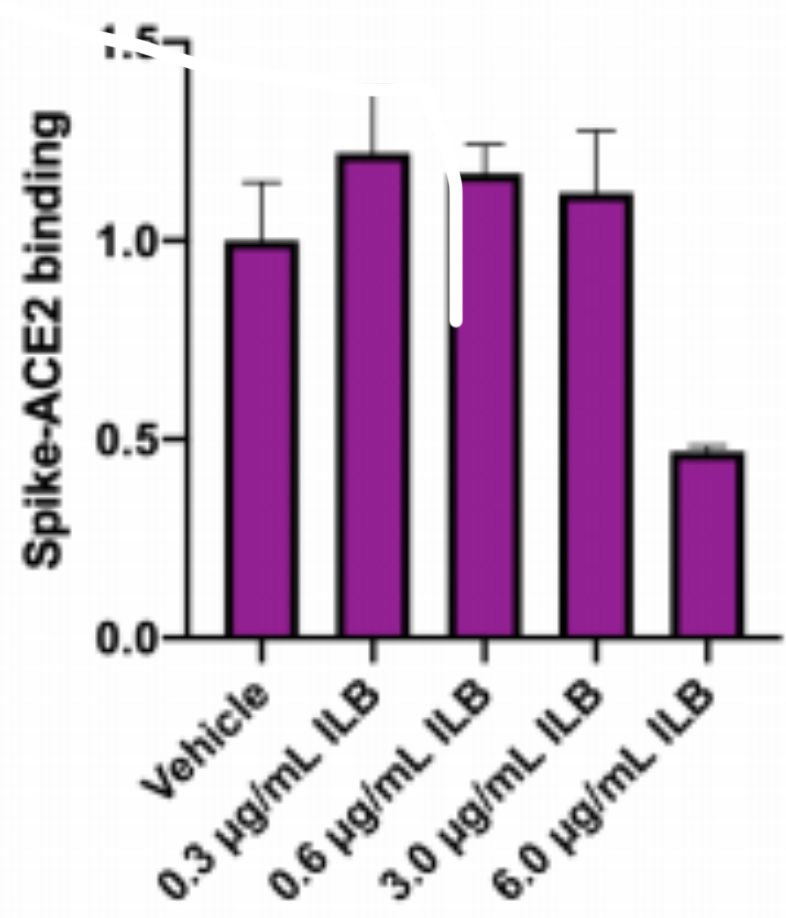
Impact of various concentrations of ILB^®^ upon SARS-CoV-2 spike protein interacting with ACE2 assessed using the RayBiotech^®^ Life ELISA assay. Each bar represents the mean data from at least 3 replicate wells.

### 3. ILB^®^ attenuation of SARS-Cov-2 variant spike protein-stimulated cytokine release from human microglia

Figures 3 and 4 show that in the absence of LPS, the ‘Omicron’ Spike protein (5.0 mg/mL, in the presence of cross-linker evoked an increase in the levels of TNFα released from microglia and this was reduced by the presence of ILB^®^ (see Donors 1, 2 and 3 at 20 h in Figure 3 and Figure 4). However, a relatively low concentration of LPS (10 ng/mL) in the presence of the ‘Wuhan’, ‘Delta’ or ‘Omicron’ spike proteins evoked a significant increase in IL-6 and TNFα release that was reduced by the presence of ILB^®^ (see Donor 1 at 20 h; Donor 2, at 6 and 20 h; Donor 3 at 20 h in Figure 3 and Figure 4), and an increase in IL-1β that was reduced by the presence of ILB^®^ (see Donor 2 at 20 h).

**Figure 3.**
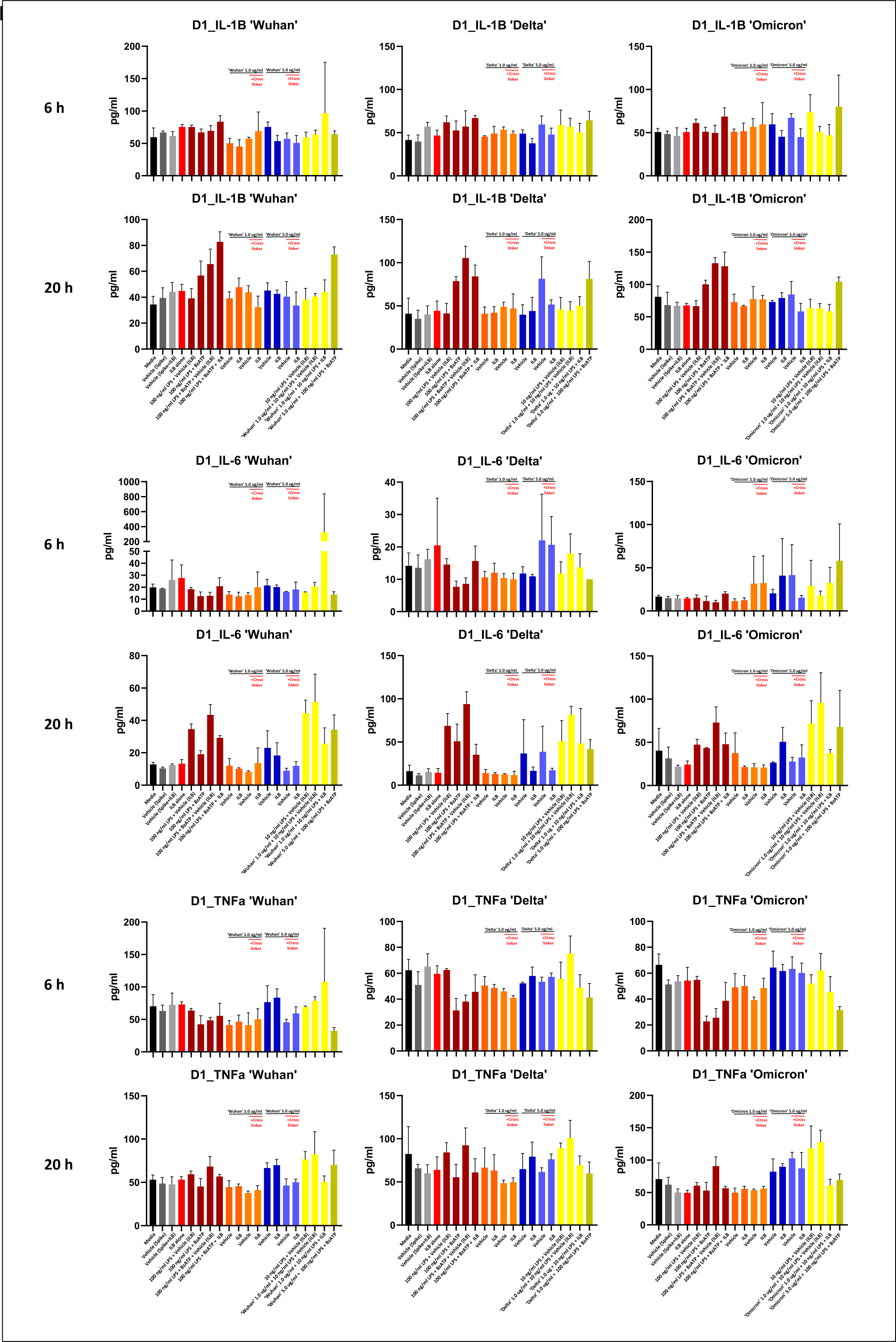
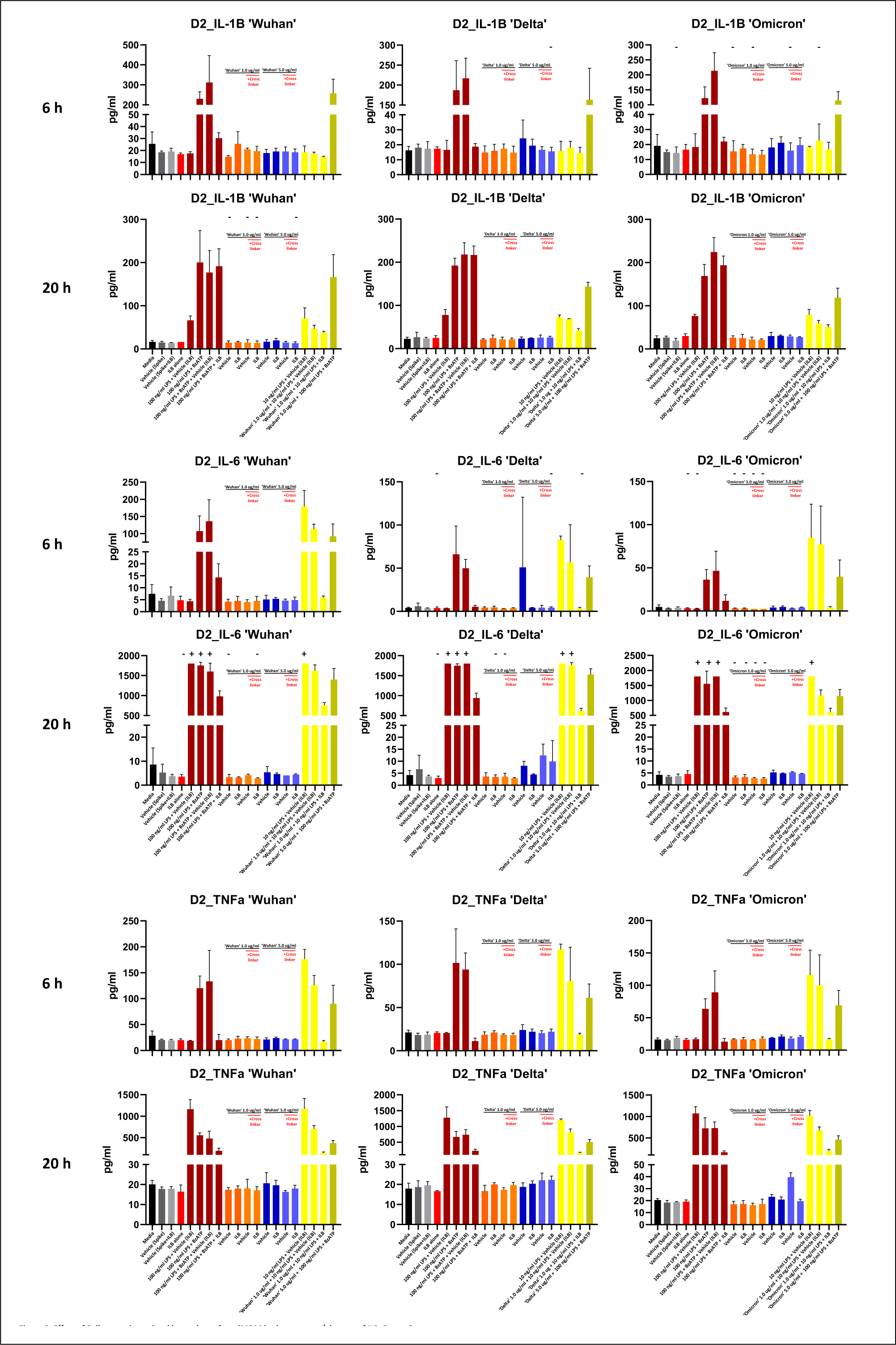
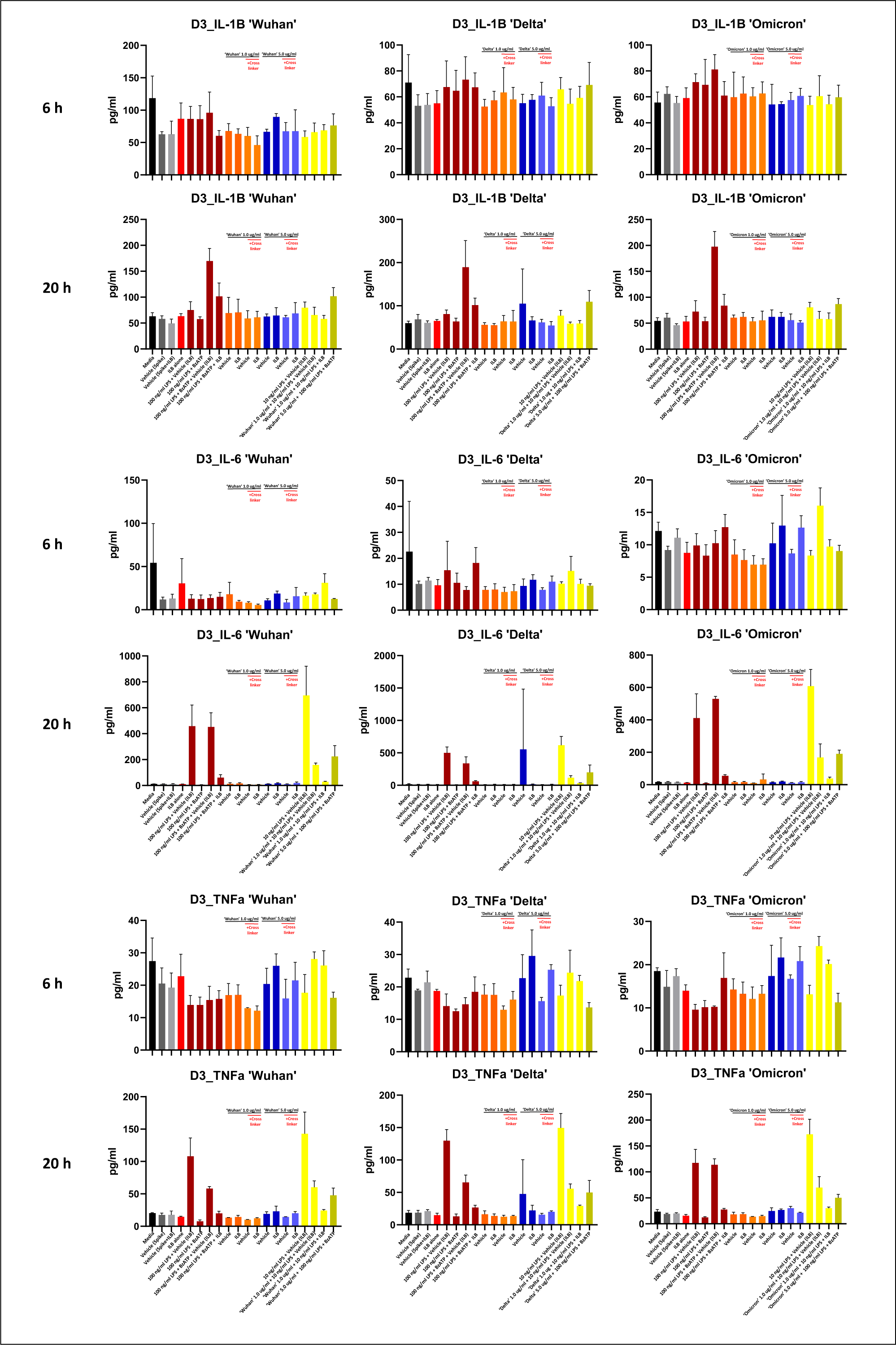
Effect of SARS-CoV-2 variant spike protein (Wuhan, Delta and Omicron) on cytokine release from iMDM in the absence or presence of ILB^®^. Data presented are mean + SD arising from triplicate cultures. Data presented is from 3 individual Donors (D1-3).

**Figure 4.**
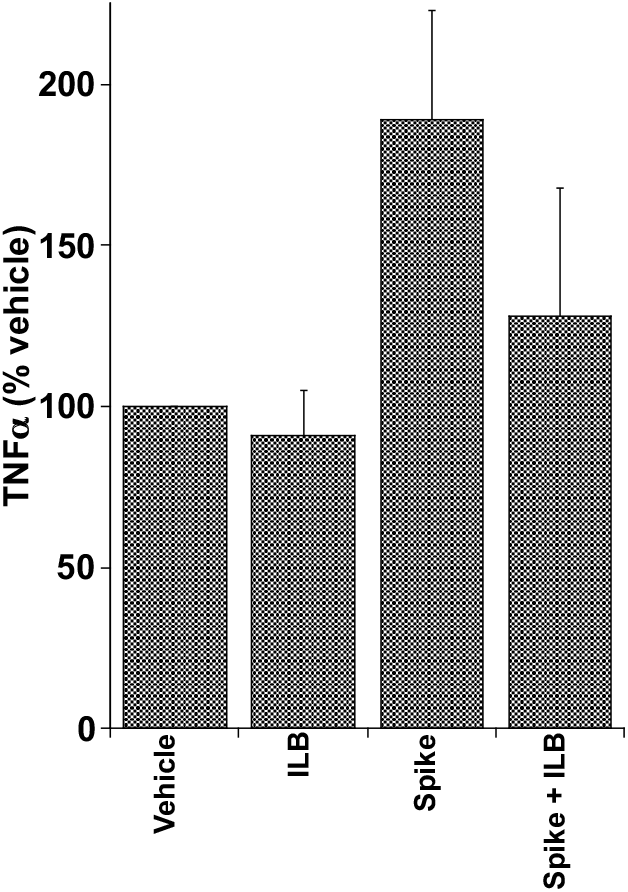
ILB^®^ attenuation of SARS-CoV-2 omicron spike protein-stimulated TNFα release from iMDM. Data presented are mean + SEM arising from cells derived from 3 individual donors (D1-3).

## Discussion

Under the conditions tested, ILB^®^ displayed neutralising activity against infection by human coronavirus NL63 when pre-incubated with the cells for 1h prior to infection, with an EC_50_ of 5.90 mg/ml. Cytotoxicity was observed only at the higher concentrations of ILB^®^, with a TC_50_ of 56.5 mg/ml. The ability of ILB^®^ to prevent cell infection is supported by the observation of a direct inhibition of the interaction between SARS-CoV-2 spike protein and ACE2. Taken together, this evidence of anti-viral activity would be predicted to be beneficial to patients by reducing the ability of the SARS-CoV-2 virus to gain entry to cells. The anti-infective activity of ILB^®^, a clinical grade drug with a proven safety profile in humans^8^, points to its substantial potential as a new adjunct anti-viral drug to help counter the uncontrolled growing global burden of Coronavirus disease.

Anti-viral drugs can be divided into two major categories: those directed against the host and those directed against the vector. Anti-viral compounds directed against host targets are of interest in the search for broad spectrum anti-viral compounds that may be able to address present or future unknown viral emergences, since they can be directed at pathways that are common to multiple virus types. Of importance, compounds that target the host pathway exploited by viruses may be expected to show a higher barrier to resistance, which is of importance for RNA viruses with a relatively high tendency to mutate, such as Coronaviruses.

Highly sulphated glycosaminoglycans (GAGs) like heparan sulphate are found in and around animal cells and are involved in the infection of many pathogenic enveloped and non-enveloped viruses, including Coronaviruses like SARS-CoV-2^9-12^. These viruses utilize cell surface glycoconjugates as cellular receptors for attachment, which enable them to take their first step toward establishing infection. The binding of these viruses to cell surface heparan sulphate could be specific but also could be due to nonspecific electrostatic interactions. Of biological relevance, either possibility suggests the application of heparan sulphate mimetics as anti-viral therapies. Indeed, soluble GAGs, especially heparin mimetics, have been used as competitive inhibitors to block Coronavirus infection^13-15^. Of relevance, the LMW-DS used in this study, ILB^®^, acts as a soluble heparin mimetic that can offer a competitive binding site for heparin-binding moieties, thereby preventing receptor interactions^16^. It therefore seems probable that, in this instance, soluble ILB^®^ is acting competitively with cell surface heparan sulphate to bind virus and block cell attachment and internalization *via* spike protein:ACE2 receptor interactions.

Extensive preclinical and clinical studies, reported by us elsewhere^8,16-19^, suggest that ILB^®^ may also be a useful tool to counteract the post-viral syndrome that is often associated with Coronavirus infection. Accordingly, ILB^®^ activates natural repair processes to control hyperinflammation and thrombosis, normalise cellular metabolism and remodel damaged tissues^16^. Thus, uniquely, ILB^®^ acts as a broad acting anti-viral drug that targets every step in the progression of viral disease from infection through to cellular pathology. This inference of restoration of cellular and tissue homeostasis in humans is further supported by the studies reported here on ILB^®^ control of cytokine release from human microglia stimulated with the omicron spike protein variant that suggest the ability of ILB^®^ to counteract chronic viral neuroinflammation.

In contrast to previous reports^20-26^, by other research groups investigating the ‘Wuhan’ SARS-CoV-2 spike protein with myeloid cells, a clear response to ‘Wuhan’ SARS-CoV-2 spike protein-evoked cytokine response (proposed to act *via* TLR4^20-24^) was not evident in our studies^20,21^ using myeloid induced-monocyte derived microglia (iMDM). LPS contamination of preparation remains a major challenge when studying TLR biology^25^, especially for bacteria derived proteins^20^ (where trace levels of LPS may remain), and whilst others used mammalian^21,22,24^ or insect^23^ cell line-derived spike protein the potential for bacterial contamination remains. Although there are differences in the source of Wuhan spike protein used in previous studies compared to the present study, the SARS-CoV-2 spike proteins used in the present study are generated using mammalian expression systems, and furthermore their direct ability to bind ACE2 has been verified using SPR analysis and also a spike protein binding assay to mammalian cells that was reversible by anti-spike protein neutralising antibodies^27^. Similar validation is evident for the delta and omicron spike proteins used in the present study. It is therefore of interest that the Wuhan and delta spike proteins did not evoke evident cytokine release from the iMDM. In contrast, the omicron spike protein (5.0 mg/mL), in the presence of cross-linker in an attempt to recapitulate the native spike presentation to the surface of the cell, enhanced consistently TNFα release from the iMDM cells and of potential therapeutic relevance, the enhanced release of TNFα was reduced consistently by ILB^®^. An ability of ILB^®^ to reduce the release of pro-inflammatory cytokines from virally ‘primed’ microglia has relevance to potential beneficial effects of ILB^®^ to treat numerous post-viral syndromes including long COVID^28-30^.

In conclusion, ILB^®^ displays potential as a host-directed anti-viral drug that inhibits coronavirus infection and may be a useful adjunct anti-viral treatment. In addition, the anti-inflammatory activity of ILB^®^ suggest this drug also has potential to alleviate some of the long-term consequences of viral infection in central and peripheral tissues. The clinical stage of development of ILB^®^ offers the opportunity to test the anti-viral and post-viral syndrome activities summarised in Figure 5 in clinical trial to determine if the results translate for the benefit of patients infected with Coronaviruses.

**Figure 5.**
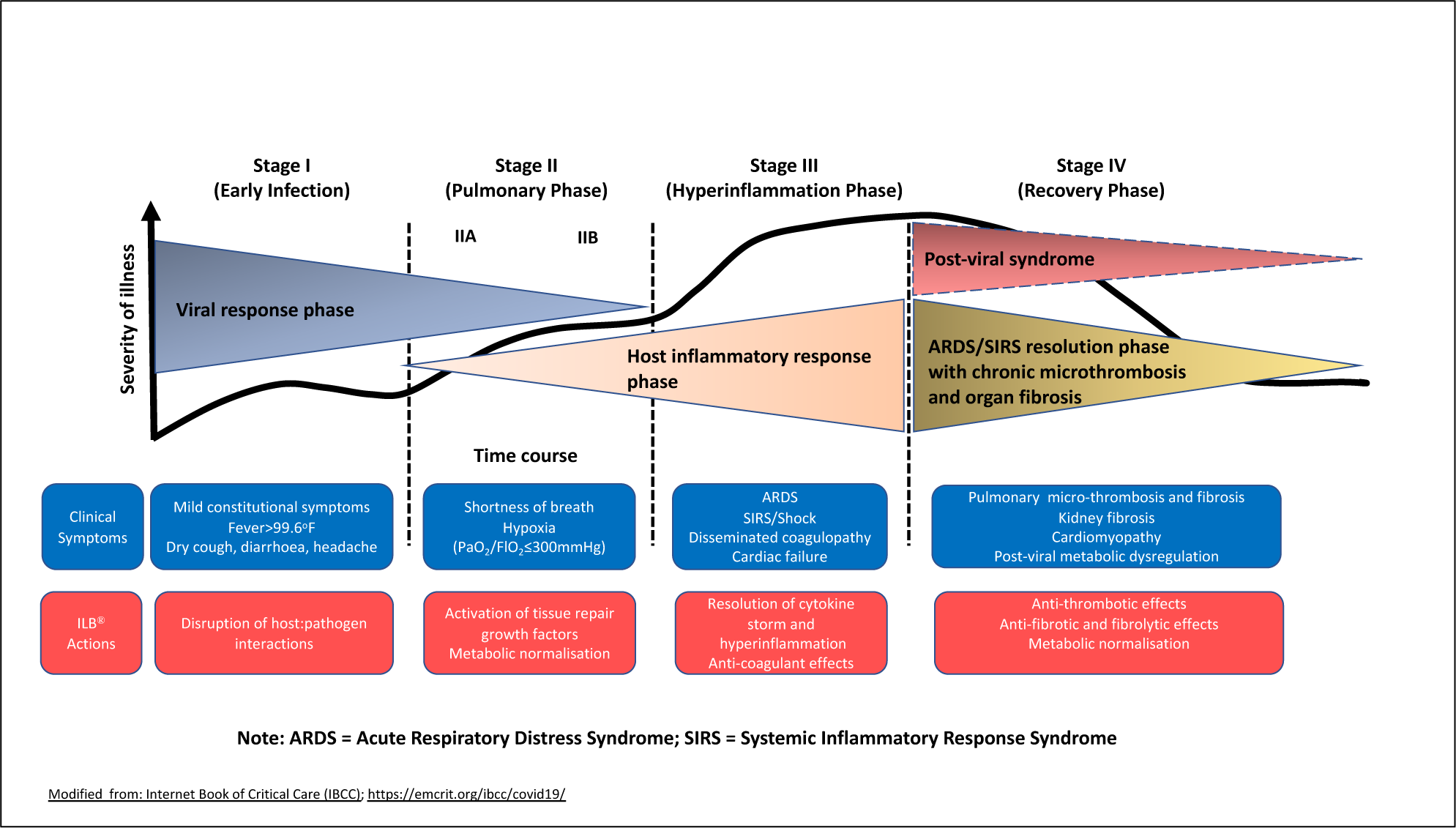
ILB^®^ activities indicating potential as an anti-viral/post-viral therapy

## Acknowledgements

This work was funded by Tikomed AB.

## Appendix: Supplementary results

### Percentages of infection – IF assay

Table S1 shows the percentages of LLC-MK2 cells infected by human coronavirus NL63 and treated with formulations of ILB^®^ or Remdesivir (assay control) at 48h post-infection. Eight dilutions were tested as indicated in the table. Three technical replicates were performed. Untreated infected and untreated uninfected controls were included (both with media and media without diluent). Wells without cells were also included.

**Table S1.**
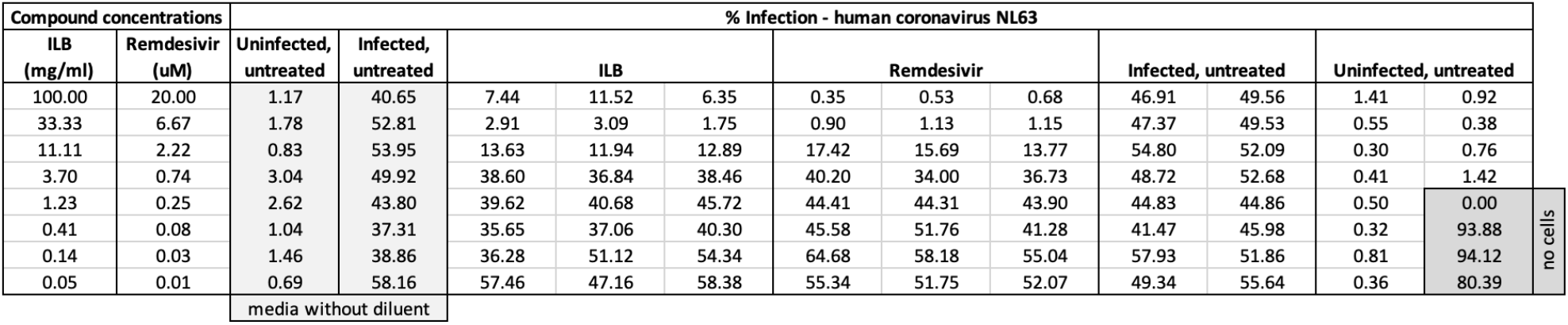
Percentages of infection at 48h.

### Percentages of cytotoxicity

Table S2 shows the percentages of cytotoxicity of LLC-MK2 cells incubated with formulations of ILB^®^ or Remdesivir (assay control) for 48h. For both compounds, eight dilutions were tested. Three technical replicates were performed. Untreated controls were included (both with media and media without diluent). No cell and Triton X-100-treated controls were also included.

**Table S2.**
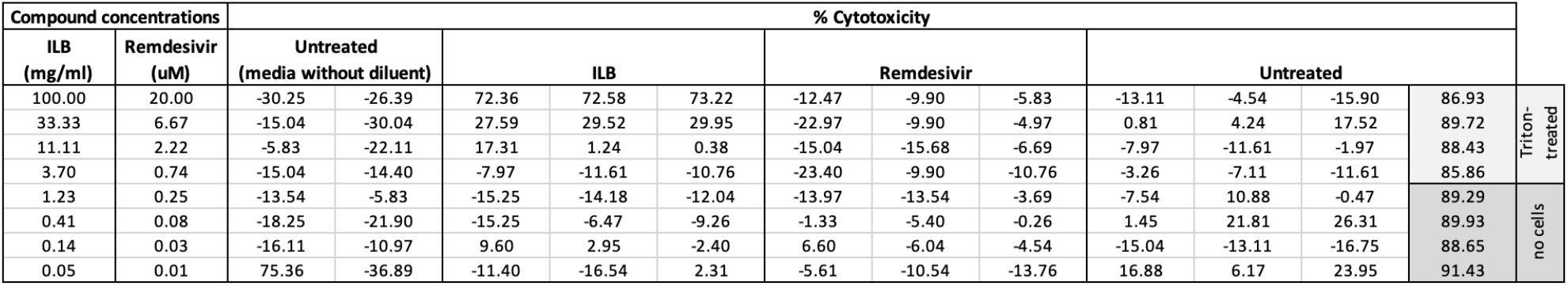
Percentages of cytotoxicity at 48h.

